# Excitation-contraction coupling and its relation to synaptic dysfunction in Drosophila

**DOI:** 10.1101/2020.07.09.196261

**Authors:** Kiel G. Ormerod, Anthony E. Scibelli, J. Troy Littleton

## Abstract

The Drosophila neuromuscular system is widely used to characterize synaptic development and function. However, little is known about how specific synaptic deficits alter neuromuscular transduction and muscle contractility that ultimately dictate behavioural output. Here we develop a system for detailed characterization of excitation-contraction coupling at Drosophila larval NMJs and demonstrate how specific synaptic and neuronal manipulations disrupt muscle contractility. Muscle contraction force increases with motoneuron stimulation frequency and duration, showing considerable plasticity between 5-40 Hz, while saturating above 50 Hz. Temperature is negatively correlated with muscle performance and enhanced at lower temperatures. A screen for modulators of muscle contractility led to the identification and characterization of the molecular and cellular pathway by which a specific FMRFa peptide, TPAEDFMRFa, increases muscle performance. These findings indicate *Drosophila* NMJs provide a robust system to relate synaptic dysfunction to alterations in excitation-contraction coupling.

## Introduction

Neuromuscular systems that regulate stereotyped motor behaviours are composed of multiple component parts that interact in a highly coordinated and synchronized fashion to control muscle contraction and displacement (Goulding, 2009; Selverston, 1980). Central pattern generators (CPGs) initiate stereotyped behaviours such as locomotion, feeding, and swimming (Katz and Frost, 1995; Schwarz et al., 2017). CPGs relay rhythmic output through interneurons to motoneurons, controlling the timing and magnitude of muscle contraction (Selverston, 2010). In turn, the peripheral nervous system (PNS) signals the dynamic state of the muscle back to the CPG to modulate circuit output (Singhania and Grueber, 2014). A large body of work on the composition and regulation of these component parts comes from invertebrate studies in crayfish, leech, locust, crab, lamprey and others (Marder et al., 2005; Marder and Bucher, 2007; Nusbaum et al., 2017). *Drosophila melanogaster* has become a popular system to dissect and explore the composition and regulation of many of these components, including CPGs (Clark et al., 2018; Song et al., 2007), interneurons (Hasegawa et al., 2016), and sensory neurons (Singhania and Grueber, 2014; Song et al., 2007). Despite these efforts, studies directly examining the role of excitation-contraction coupling in locomotion are limited (Lehmann and Dickinson, 1997; Ormerod et al., 2018, 2016, 2015).

Locomotion is one of the most well studied rhythmic behaviours, including in Drosophila (Caldwell et al., 2003; Cheng et al., 2010; Clark et al., 2018; Hughes and Thomas, 2007; Saraswati et al., 2004; Song et al., 2007). Drosophila larval locomotion encompasses different types of movements (e.g. turning, rolling, burrowing) but motivated linear crawling is the most highly stereotyped and orchestrated repetitive patterned motor behaviour produced from identical contraction waves (Heckscher et al., 2012; Hughes and Thomas, 2007). Muscles directly below the cuticle contract, then relax, to propagate peristaltic contraction waves from the posterior to the anterior of the larvae. These peristaltic waves are achieved by systematic, coordinated contractions of bodywall muscles, predominately those in the abdominal segments (Clark et al, 2018). *Drosophila* larvae are bilaterally symmetrical with stereotyped abdominal hemisegments composed of 30 individual muscle fibers. Each muscle is a viscerally-located, supercontractile (contracting to a length less than 50% of resting length) striated multinucleated muscle fiber (Keshishian et al., 1996) attached directly to the cuticle through apodemes (Koh et al., 2000).

Canonical muscle fiber contractility is controlled via postsynaptic glutamate receptors activated by glutamate released from presynaptic motoneuron terminals (Harris and Littleton, 2015). Approximately 36 motoneurons per hemisegment innervate the abdominal musculature. Three different motoneuron subtypes are present: type I, which releases glutamate from synaptic vesicles; type II, which predominantly contain large dense core vesicles (DCVs); and type III, which innervate muscle 12 and release insulin-like peptides (Budnik, 1996; Hoang and Chiba, 2001). The neuromuscular junction (NMJ) is a critical site of plasticity and can be controlled by neuromodulators released directly from motoneuron terminals or centrally into the open circulator system as hormones (Milakovic et al., 2014; Ormerod et al., 2018, 2013). Type I motor neurons are rhythmically active during waves of muscle contractions underlying forward and reverse locomotion (Newman et al., 2017). Motoneuron activity is dictated by the neuronal input received from descending interneurons, whose firing pattern is ultimately controlled by the locomotor CPGs within the ventral nerve cord (VNC) (Fox et al., 2006).

To dissect the relative contribution of excitation-contraction coupling on locomotion, we employed a force transducer with 10 μN resolution and a unique motoneuron-stimulation paradigm to characterize the dynamic output of the neuromuscular system and how plasticity in synaptic function manifests in changes to muscle output. Synaptically-driven muscle force increases with stimulation frequency and duration, with increases in either component leading to increased force. However, increasing both factors ultimately causes saturation in muscle force. The most substantial plasticity in muscle contractility occurs between 5 and 40 Hz stimulation. We also examined how mutants in several critical synaptic proteins, including Synaptotagmin (SYT), Complexin (CPX), and Glass-bottom boat (GBB), alter muscle performance. In addition, we found that temperature has a critical effect on *Drosophila* muscle contraction. Using this force contraction assay, we screened potential modulators of muscle contractility and identified the neuropeptide, TPAEDFMRFa, that increases muscle performance in a dose-dependent manner. We characterized the effects of the FMRFa peptide on muscle contractility and defined components of the pathway though which it operates. These data directly relate synaptic function with excitation-contraction coupling, providing insights into how key synaptic proteins control not only neurotransmitter release and synapse development, but also how they contribute to muscle contractility.

## Results

To explore the contractile properties of Drosophila 3^rd^ instar larval NMJs, muscle contractions were elicited in semi-intact preparations with the CNS removed. A force transducer with 10 μN resolution was modified to attach to the posterior end of a larvae (Figure 1A). Previous studies in Drosophila examined contractile properties by continually stimulating motoneurons at the same intraburst frequency and duration (Ormerod et al., 2016). An example of a single contraction induced from the setup, highlighting the amplitude, rise tau, and decay tau is shown in Figure 1B. To systematically characterize muscle contraction performance, bodywall contractions were induced by stimulating motoneurons at a frequency within the physiological range of 40 Hz for 600 ms duration every 15 s (0.067 Hz) for up to an hour. Muscle responses were robust with steady-state contractions recorded after 1 hour showing a 29.2 + 9.6% reduction in force over this period (Figure 1C, N=10). Endogenous muscle contractions are driven by neuronal input whose firing pattern is ultimately controlled by the locomotor CPGs within the ventral nerve cord (Song et al, 2007). We recorded fictive locomotor patterns within motoneurons encoded from the CPGs from bodywall muscles via intracellular recordings from semi-intact preparations with the CNS and VNC left intact (Figure 1D). These patterned outputs displayed widely varying intraburst frequencies from 1 to 150 Hz. Given this variable endogenous activity, we explored a more dynamic approach to eliciting muscle contractions to determine the range of muscle force bodywall muscles are capable of producing under a variety of motoneuron stimulation frequencies.

**Figure 1.**
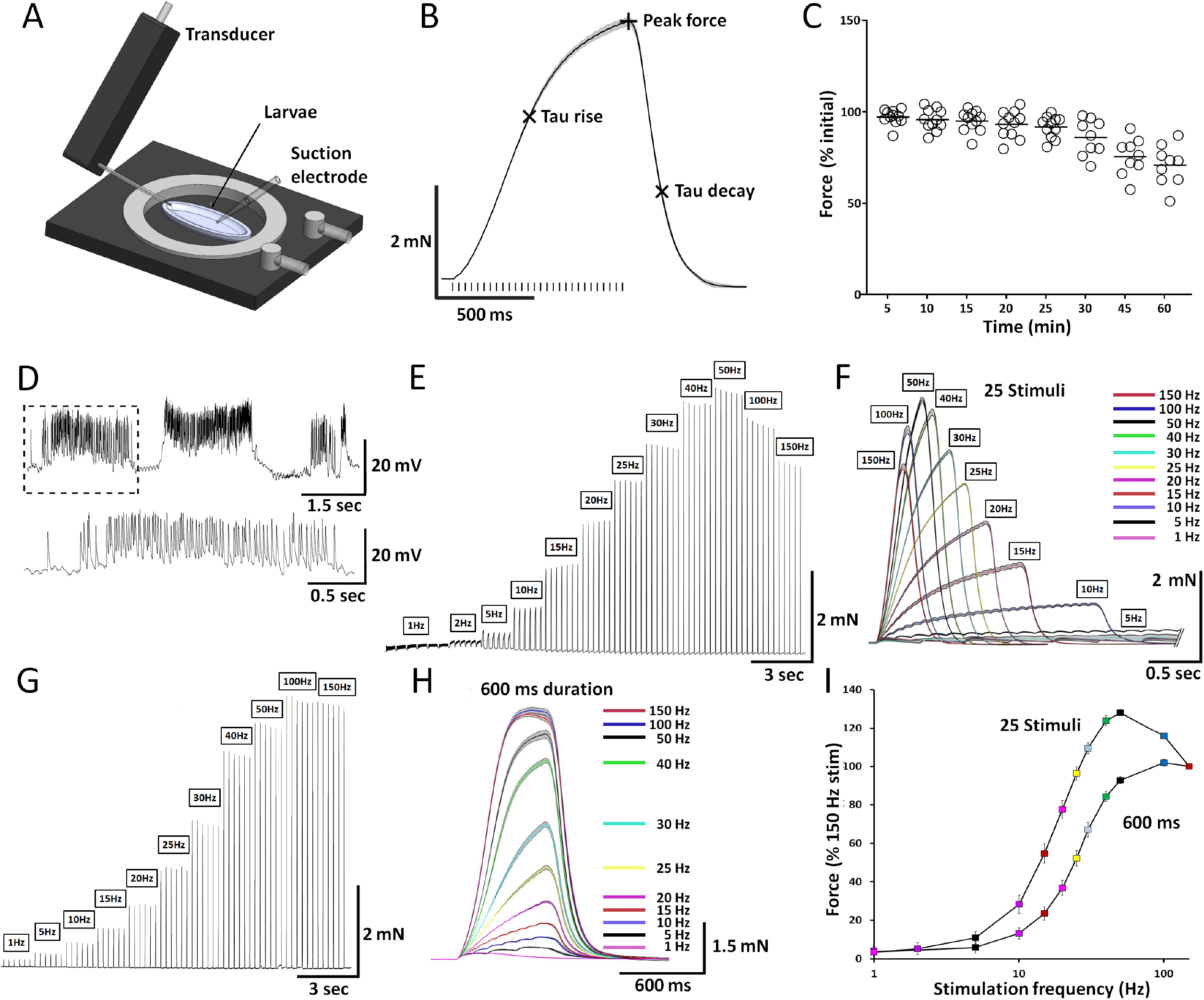
Muscle contraction dynamics from 3^rd^ instar larvae. **A)** 3D schematic of the experimental setup highlighting the force transducer, temperature-controlled holder with perfusion system, suction electrode, and 3^rd^ instar larva. **B)** Representative trace from a single contraction elicited at 40 Hz for 750 ms. Crosshairs indicate metrics derived from each contraction used for subsequent analyses. **C)** Percent of initial contraction force for steady-state recordings from control Canton S larvae elicited with 40 Hz stimulation for 600 ms every 15 s. Circles denote individual replicate recordings plotted as a percentage change in contraction amplitude compared to the initial contraction. N=9-11. **D)** Representative intracellular voltage recording from muscle fiber 6 demonstrating CPG firing patterns from a larva with the CNS left intact. The dashed box in the upper trace is expanded below. **E)** Larval motoneuron stimulation paradigm used to generate dynamic force-frequency recordings for muscle contraction from initial threshold through saturation (1-150 Hz). Stimulus number was kept constant at 25. **F)** Six replicate contractions from each stimulation frequency in panel E were averaged and plotted with the associated 95% CI. **G)** Results using dynamic force-frequency motoneuron stimulation paradigm where stimulus duration was kept constant at 600 ms **H)** 6 replicate contractions from each stimulation frequency in panel G were averaged and plotted with 95% CI. **I)** Muscle force versus stimulation frequency from each of the stimulation paradigms depicted in E-H is shown (25 stimuli: intraburst duration consisted of 25 stimuli at each stimulus frequency, N=10, 6 replicate contractions were elicited at each stimulation frequency and averaged for each animal. 600 ms duration: intraburst duration at each frequency was 600 ms in duration, 6 replicate contractions were elicited at each stimulation frequency and averaged for each animal, N=20).

Initially, 25 impulses were delivered to motoneurons and the stimulation frequency was varied from 1 to 150 Hz. For each experiment, 6 replicate contractions were induced at each stimulation frequency, followed by the next stimulation frequency, et cetera (Figure 1E). Contractions were averaged across the 6 replicate stimuli for a given stimulation frequency and the resulting trace with 95% confidence interval (CI) was determined (Figure 1F). A force-frequency curve of the force generated from this stimulation paradigm was then plotted as a percentage of the magnitude of force generated at the highest stimulation frequency of 150 Hz (Figure 1I, N=8). 25 stimuli at 1 Hz stimulation induced 3.5 + 1.1% of the maximum force produced at 150 Hz, while 2 Hz stimulation was sufficient to induce 5.2 + 1.1% of the maximum force. Force generation at 2 Hz was not significantly greater than 1 Hz stimulation. In contrast, 25 stimuli at 5 Hz produced a resultant force of 11.0 + 1.1% of the force at 150 Hz, indicating this stimulation frequency is sufficient for temporal summation for contractile force. From 10 to 50 Hz stimulation, a nearly linear increase in force was produced at each successive increase in frequency (10, 15, 20, 25, 30, 40, 50 Hz), producing a maximal force at 50 Hz. Increasing the stimulation frequency beyond 50 Hz resulted in a reduction in force production. Therefore, when total stimuli number remains constant, increasing stimulation frequency increases force production until saturation is reached at 50 Hz.

Although 50 Hz drove maximal muscle contraction, it was unclear whether this saturation of force generation was a consequence of the stimulus frequency or stimulus duration. To explore this relationship further, burst duration was held constant at 600 ms and the frequency of stimulation was varied from 1 to 150 Hz. Figure 1G depicts the raw data from each individual contraction within a single trial, and Figure 1F depicts averaged traces from the replicate stimuli at a given stimulation frequency with a corresponding 95% CI. Under these conditions, 5 Hz stimulation produced a force of 6.0 + 3.1% of that produced during 150 Hz stimulation (Table 1). Comparable to the previous experiment, force production increased in a linear fashion as the frequency increased from 10-50 Hz. However, increasing stimulation frequency to 100 and 150 Hz increased the force. Thus, both the duration and frequency of stimulation are critical factors in determining the absolute magnitude of force production.

**Table 1:**
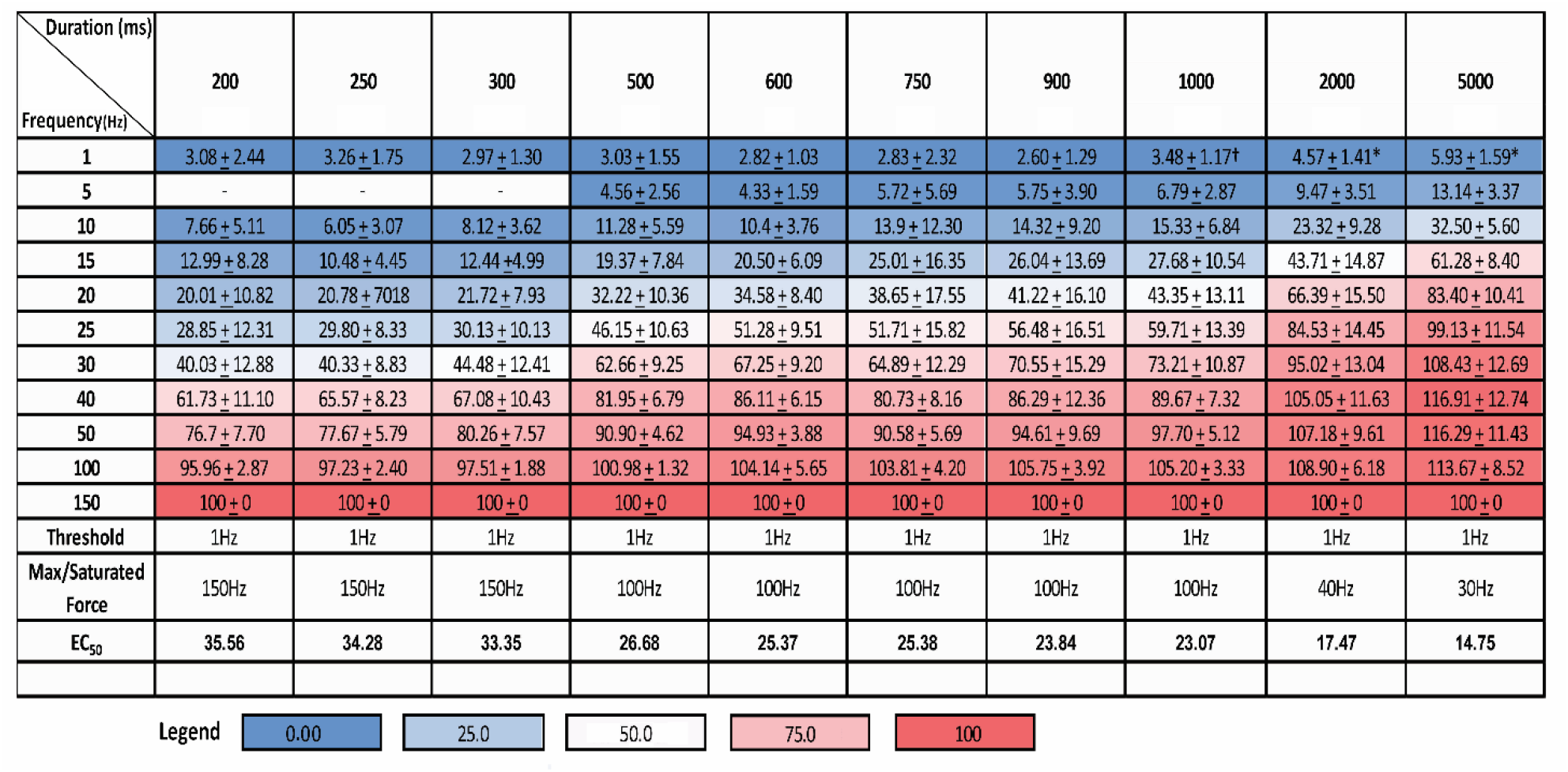
Force-frequency values for 10 different stimulus durations calculated as a percentage of the contraction force elicited from 150 Hz stimulation. Thresholds indicate the stimulation frequency needed to elicit a visible and quantifiable contraction trace. Maximum or saturation force value indicates the frequency required to elicit a contraction that is equal in magnitude or exceeds the contraction force value obtained using 150 Hz stimulation. The EC_50_ denotes the stimulation frequency required to generate a contraction that is 50% of maximal force value obtained using 150 Hz, calculated from the force-frequency plots using GraphPad Prism.

To more fully explore the relationship between stimulus frequency and stimulus duration, we generated a series of force-frequency curves by keeping the burst duration constant through an entire experiment over 200, 250, 300, 500, 600, 750, 900, 1000, 2000, or 5000 ms while increasing stimulation frequency from 1 to 150 Hz. Each of the independent force-frequency curves for a given burst duration generated a sigmoidal curve similar to what was observed at 600 ms (Figure 2A and Table 1). The overall patterns from the stimulation force recordings revealed that increasing stimulus frequency or duration results in a progressive and gradual increase in muscle force production. In all experiments, a single action potential was sufficient to induce a contraction that was between 2.6 and 3.3% of the 150 Hz contraction force (note that 1 Hz stimulation at 1000, 2000, and 5000 ms corresponded to 2, 4, and 10 stimuli respectively). Stimulus durations of 200, 250, and 300 ms resulted in increased force production with increases in stimulus frequency that did not saturate until 150 Hz (Table 1). Stimulus durations of 600, 750, 900, and 1000 ms resulted in sigmoidal increases in force production with increases in stimulus frequency that saturated at 100 Hz. Increasing the stimulus duration to 2000 and 5000 ms resulted in a saturation of force at 40 and 30 Hz, respectively. Given force saturates at or above 100 Hz for all stimulus durations, there should be no differences between these conditions once force reaches maximal. Thus, it is not surprising that the greatest differences between stimulus durations are seen from 10 to 50 Hz, where considerable plasticity in muscle performance is still possible (Fig 2A). Plotting the percent differences between the various stimulation durations as a function of stimulus frequencies generated a bell-shaped curve with peak differences observed at 25 Hz. The greatest effect of stimulus duration was observed at 25 Hz, where an increase from 200 to 5000 ms resulted in a 70.3% increase in muscle force. Figure 2B depicts an overlay of 25 Hz stimulation from 200, 300, 600, 900, and 2000 ms. A critical feature of the dataset is reflected in the stimulation frequency required to generate half-maximal force (50%) for each stimulus duration, where a linear decrease in the stimulus frequency required to reach 50% is observed (Table 1). Plotting the 50% max value as a function of stimulus duration generated a strong negative correlation (R^2^=0.9) with a slope of −0.016, indicating that every 100 ms increase in duration shifts this value to the left by 1.6 Hz. Despite the dramatic effects that stimulation duration has on force production below 50 Hz, the saturated or maximum raw force generated from a 200 ms duration stimulus was not significant for any of the longer stimulus durations (Figure 2C, D). These findings indicate maximal force from Drosophila bodywall muscles can be generated by a 200 ms stimulus delivered at 100 Hz.

**Figure 2.**
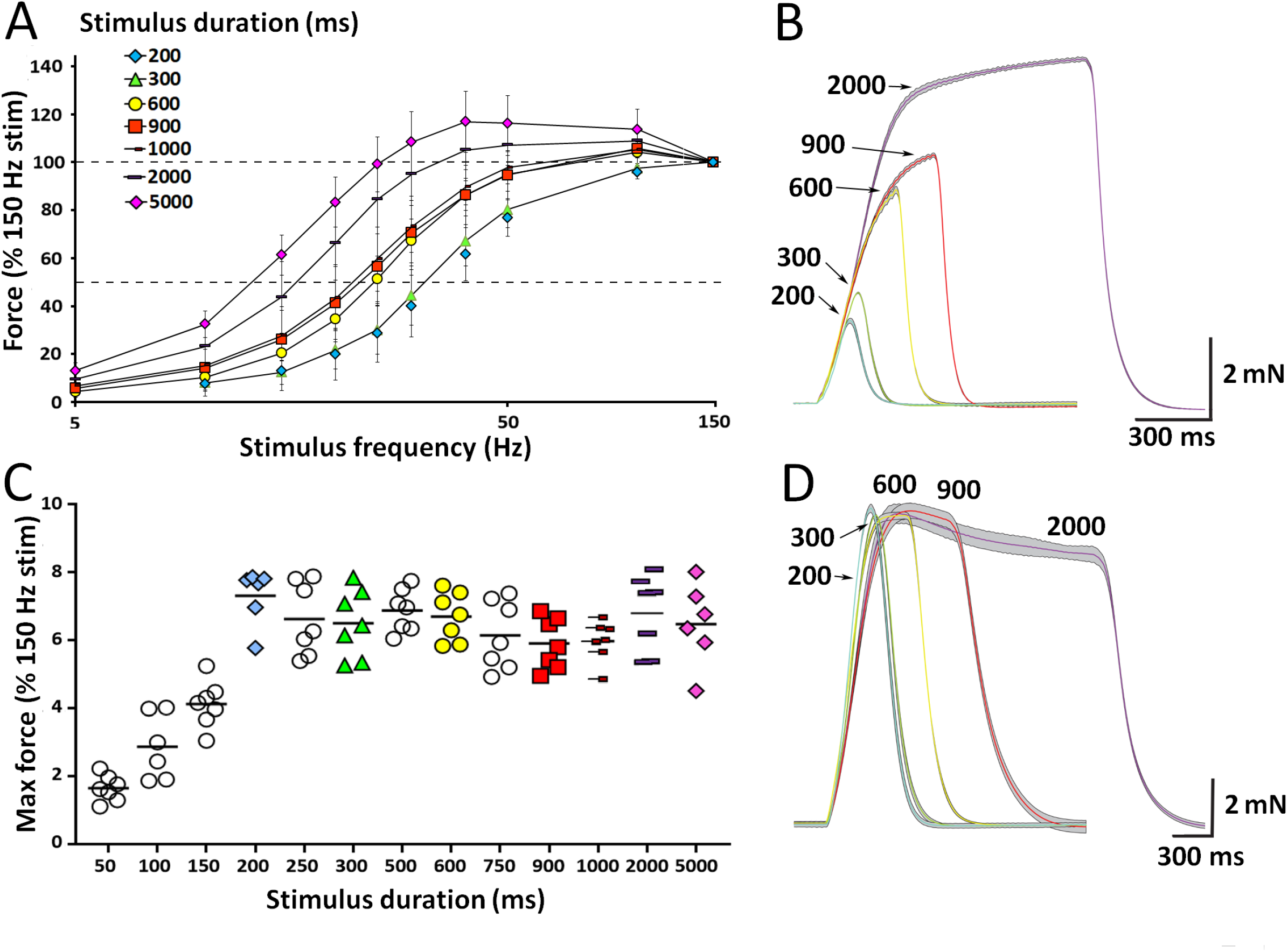
Effects of stimulus duration or frequency on muscle contraction. **A)** Cumulative force-frequency plots showing the effects of varying the stimulus duration from 200 ms to 5000 ms (N=7-10). **B)** Representative traces for 25 Hz stimulation for 200, 300, 600, 900, and 2000 ms stimulus duration. **C)** Maximal force generated from a 150 Hz stimulation frequency from each replicate larva for each of the 13 different stimulation durations. **D)** Representative single contraction traces from 200, 300, 600, 900, and 2000 ms duration stimuli at 150 Hz.

Throughout the experiments there was variability in the total raw force generated from one preparation to the next (Fig 2C), differing in magnitude by up to 50%. Given this variability, one potential contributor to the differences in force generation was larval size. To examine this variable, the length and width of 100 larvae was measured and the total force generated by each animal was determined by eliciting a 100 Hz stimulus for 600 ms (Figure 3). The larval force was then plotted against length, width, area, and volume to determine if these physical parameters correlated with overall force production. None of these metrics revealed a positive correlation with contractile force (Figure 3C, D), indicating larval size does not have a significant effect on the total force that is generated from 3^rd^ instar larval preparations (two-tailed Pearson’s correlation; Width: r=0.01462, P=0.8906; length: r=-0.1246, P=0.2391).

**Figure 3.**
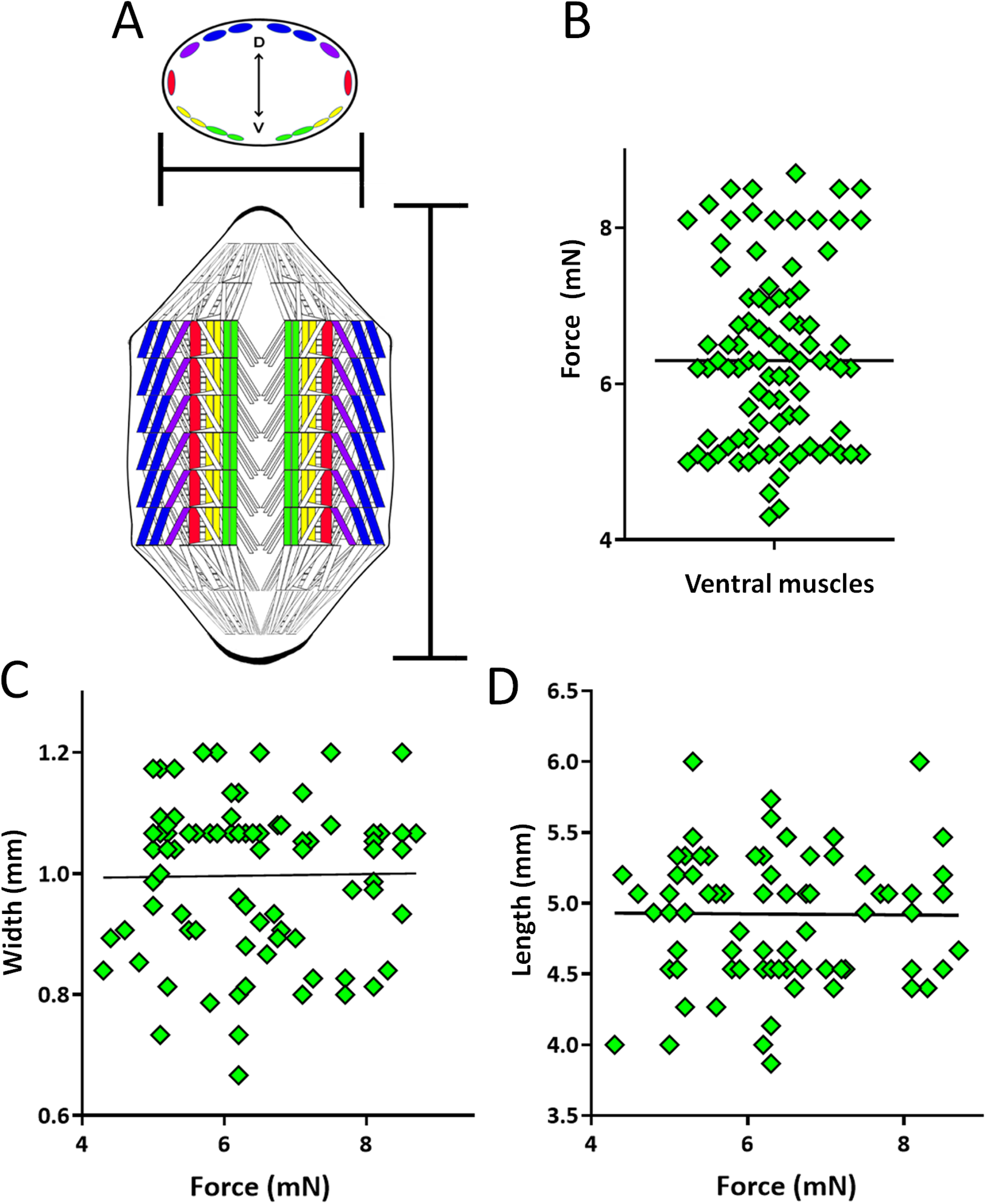
Muscle force generation is independent of 3^rd^ instar larval size. **A top)** Schematic representation of a cross-section through a 3^rd^ instar larval abdominal segment highlighting the main muscles contributing to longitudinal larval peristalsis. D-V represents the dorsal-ventral axis. Scale bar below indicates how width measurements were taken. **A bottom)** Schematic representation of a dissected larva highlighting the main longitudinal muscles that contributes to larval peristalsis. Scale bar to the right indicates how length measurements were determined. **B)** Summary of maximal force values obtained from 100 animals dissected along the dorsal midline. **C-D)** Width and length measurements and the corresponding maximal force values obtained from 100 animals. Neither parameter correlates with maximal force generated (two-tailed Pearson’s correlation; Width: r=0.01462, P=0.8906; length: r=-0.009, P=0.9325).

We next explored whether the recruitment of different muscle fibers altered muscle performance or maximal force generation. To examine the contribution from distinct muscle groups, larvae were dissected in three different orientations: i) along the dorsal midline where ventral muscle fibers 6, 7, 12, and 13 contribute predominately to longitudinal force production; ii) along the ventral midline where dorsal fibers 1, 2, 3, and 4 contribute principally; or iii) a lateral incision where lateral fibers 3, 4, 12, and 13 contribute primarily (Figure 4A, B). All three larval orientations displayed nearly identical force-frequency curves (Figure 4C, D), indicating performance was not significantly different across distinct muscle groups (One-way ANOVA, N=8, P>0.05). In addition, maximum force generated from muscle contraction elicited by stimulation in each larval orientation was similar (Figure 4E, One-way ANOVA, N=8, P=0.17).

**Figure 4.**
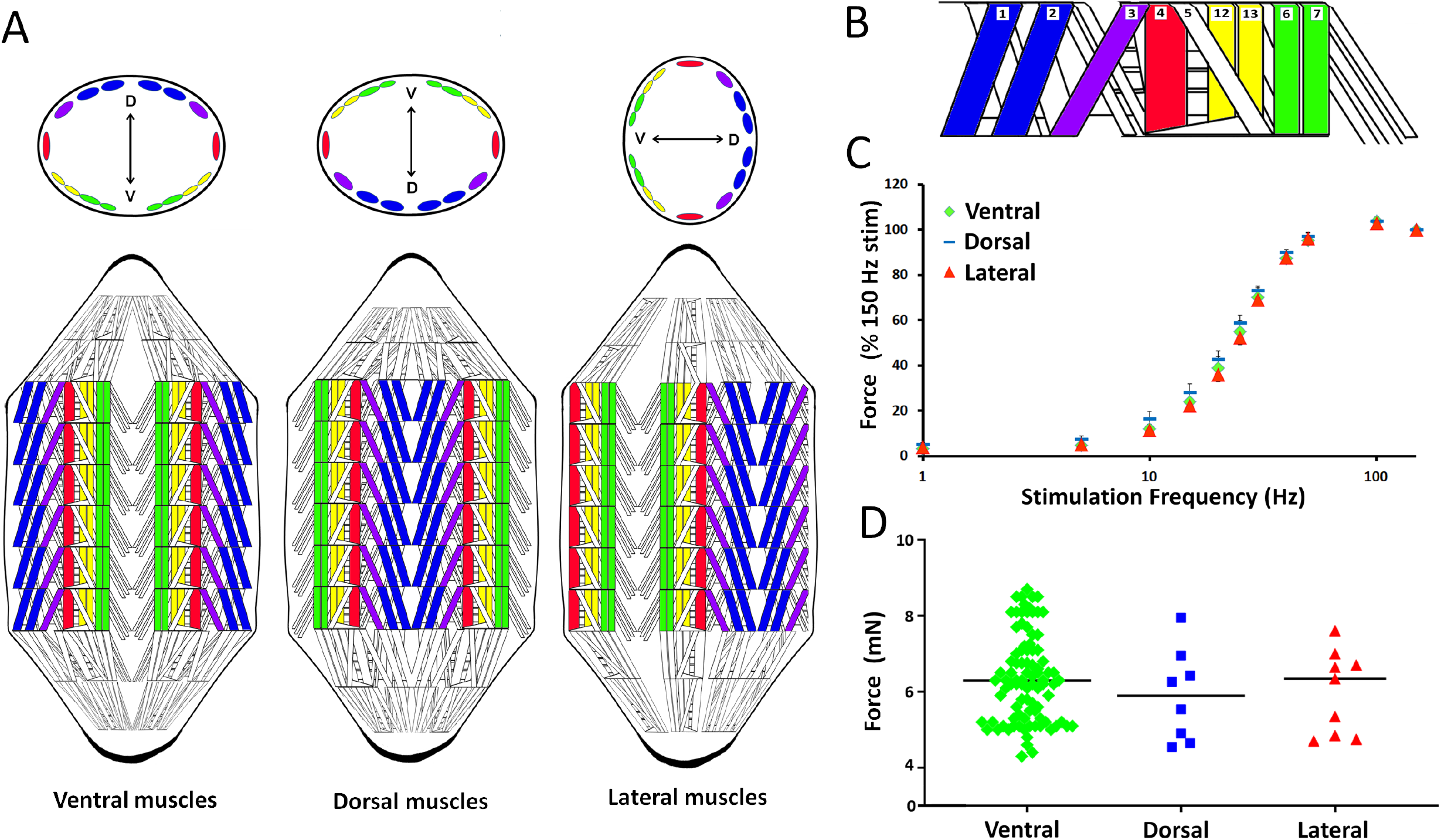
Similar contraction properties of ventral, dorsal and lateral muscle groups. **A) Top:** Schematic representation of cross-sections through larvae from three different orientations highlighting the dominant muscles contributing to force measurements in each condition. **Bottom:** Dissected larvae and the corresponding muscle configuration for each orientation. Blue denotes muscles along the dorsal axis, green denotes muscles along the ventral axis, and red denotes muscles along the medial axis. **B)** Cartoon depicting a single abdominal hemisegment highlighting muscle fiber number. **C)** Muscle force-frequency curves for each of the larval dissection orientations, indicating muscle contraction force-frequency is similar across the different fibers. **D)** Maximal force generated from a 150 Hz stimulus at 600 ms duration in each of the three different orientations shows no significant differences (One-way ANOVA, N=8, P=0.4037, F=0.915).

It is well established that both external Ca^2+^ and Mg^2+^ have a profound effect on pre-and postsynaptic intracellular mechanisms that contribute to muscle force production (Jan and Jan, 1976). Two commonly used *Drosophila* salines, hemolymph-like saline 3 and 3.1 (HL3 and HL3.1, Macleod et al, 2002) which contain 20 mM and 4 mM external Mg^2+^ respectively, were used to determine the effects of Mg^2+^ on force production. Additionally, 7 different external calcium concentrations ([Ca^2+^]_o_) were explored for each saline, along with 4 different force-frequency stimulations durations (200, 300, 600, and 900 ms, Figure 5). At 0.1 mM [Ca^2+^]_o_, the highest frequency stimulation (150 Hz) at the longest examined duration (900 ms) was unable to induce a recordable contraction in HL3. In contrast, a detectable contraction was observed at 10 Hz stimulation in HL3.1 (Figure 5A). Omitting external [Ca^2+^] resulted in no muscle contraction in either saline, even at the highest stimulation frequencies. Raising [Ca^2+^]_o_ from 0.1 to 0.25 mM was sufficient to induce contractions at higher stimulation frequencies and durations (40-150 Hz) in HL3 saline, although a profound difference remained between the two salines (Figure 5A). This force-frequency gap was also observed at 0.5 mM [Ca^2+^]_o_, and contractions were inducible at even lower stimulation frequencies in HL3.1 (Figure 5A). At 1.0 mM [Ca^2+^]_o_, no statistical differences were observed between the two salines at 600 and 900 ms duration force-frequency curves, and only subtle differences were observed at 200 and 300 ms (Figure 5A). Increasing [Ca^2+^]_o_ further to 1.5 or 2.0 mM resulted in no observable statistical differences between the two conditions. Figure 5B shows the effect of external Ca^2+^ and Mg^2+^ on the half-maximal (50%) stimulation frequency across the entire data set. Plotting the maximal force generated at each [Ca^2+^]_o_ for each stimulus duration generated data similar to that observed in Table 1, where an increase in either factor resulted in enhanced force generation in HL3. Above 1 mM [Ca^2+^]_o_, neither stimulus duration or increasing [Ca^2+^]_o_ affected maximal force, indicating saturation at these levels. A similar effect was observed for HL3.1, but only at 0.1 and 0.25 mM [Ca^2+^]_o_, as muscle force plateaued across all stimulus durations at and above 0.5 mM [Ca]_o_. There was significant differences between the two salines at each stimulus frequency and 0.1, 0.25, and 0.5 mM [Ca^2+^]_o_ between 15 and 50 Hz. However, no significant differences between the total force generated was found between the two salines above 1.0 mM [Ca^2+^]_o_. As such, the higher Mg^2+^ concentrations found in HL3 saline resulted in a robust reduction in muscle contractile force, consistent with elevated Mg^2+^ reducing Ca^2+^ entry into the presynaptic terminal.

**Figure 5.**
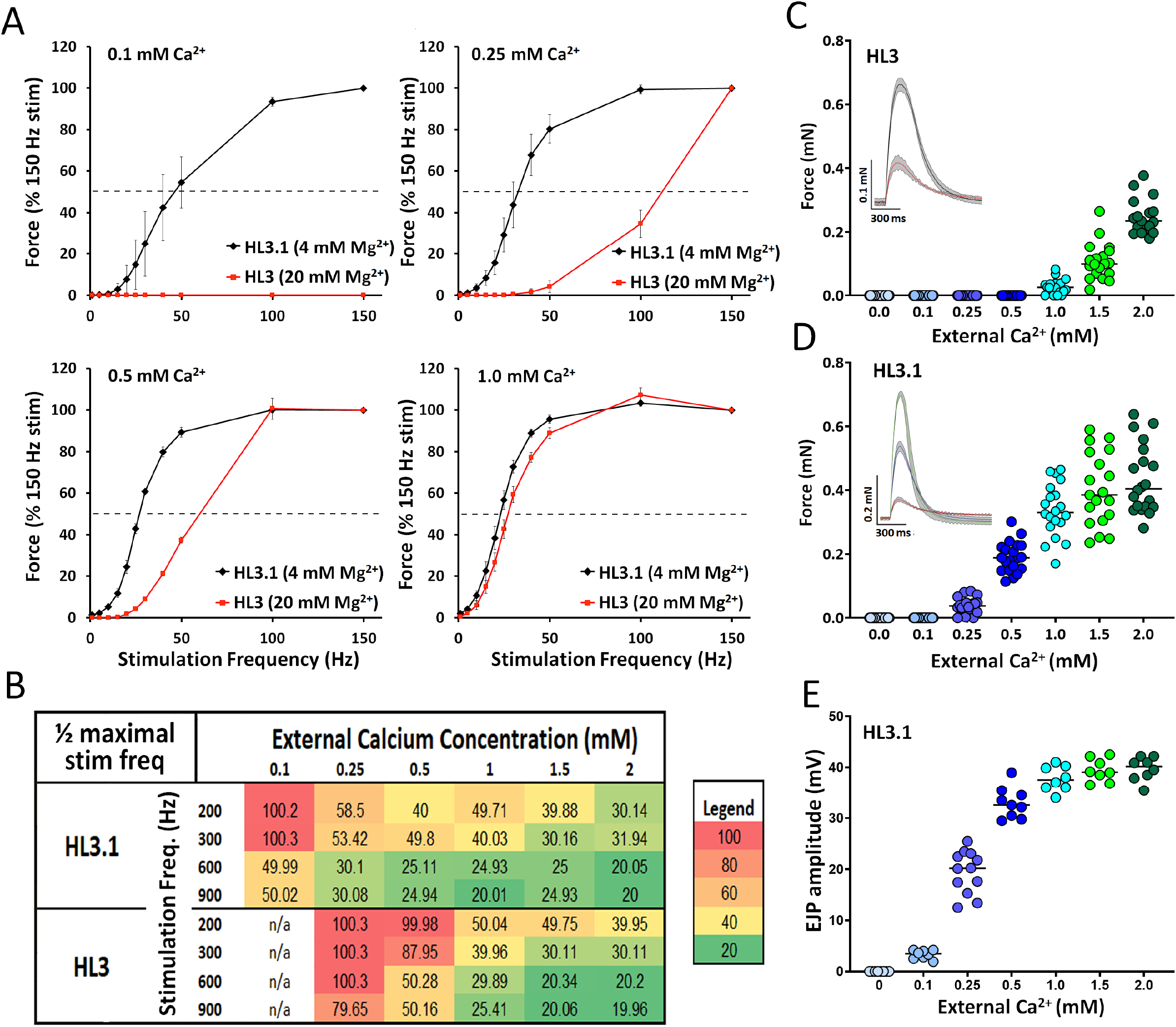
Effects of external Ca^2+^ and Mg^2+^ on muscle contraction force. **A)** Force-frequency curves 2+ generated using a 600 ms duration stimulus for two physiological salines, HL3 (20 mM Mg^2+^) and HL3.1 (4 mM Mg^2+^) in 0.1 mM [Ca^2+^]_o_, 0.25 mM [Ca^2+^]_o_, 0.5 mM [Ca^2+^]_o_, and 1.0 mM [Ca^2+^]_o_. **B)** Half-maximal stimulation frequency required to generate 50% of the maximal force observed for each stimulus duration, [Ca^2+^]_o_, and saline is shown. The data was obtained from force frequency curves generated using 200, 300, 600, and 900 ms stimulation duration for both physiological salines in 6 different [Ca^2+^]_o_. **C** Force values generated from single motoneuron stimuli in 7 different [Ca^2+^]_o_ in) HL3, and. Inset: representative force traces from 1.0 and 2.0 mM [Ca^2+^]_o_. **D** Force values generated from single motoneuron stimuli in 7 different [Ca^2+^]_o_ in HL3.1. Inset: representative force traces from 0.25, 0.5, and 2.0 mM [Ca^2+^]_o_**. E** EJPs amplitude values obtained from the 7 different [Ca^2+^]_o_ in HL3.1.

To determine if the magnitude of contractions elicited from a single stimulus represented a sensitive read-out of synaptic activity, we compared data obtained from single stimuli to that from stronger stimulations. The size of single-stimulus induced contractions as a percentage of force at 150 Hz stimulation was unchanged across stimulus durations (Table 1). The amplitude of contractions induced from a single stimulus increased in a proportional manner with increases in external Ca^2+^ and saturated beyond 1.5 mM in HL3.1 saline. While contractions elicited from single stimuli were not observed in HL3 saline until 1.0 mM [Ca^2+^]_o_, they increased proportionally from 1.0 to 2.0 mM [Ca^2+^]_o_ (Figure 5C-D). Similar proportionate increase in muscle excitability was observed in EJP recordings at the NMJ in HL3.1 saline (Figure 5E). These findings indicate contractions elicited from single stimuli, in addition to maximal force output, represent a robust mechanism to correlate synaptic function with the excitation-contraction coupling machinery.

To examine the effect of temperature on muscle performance in control larvae, contractions were generated using 600 ms duration bursts at 40 Hz every 15 seconds and plotted as a percentage of the force generated at room temperature (22°C). Surprisingly, decreasing the temperature from 22°C to 16°C increased contraction force, with a peak increase of 29.4 + 4.9% observed at 17°C (Figure 6A). Increasing the temperature from 22°C to 39°C gradually decreased the force of contraction until a complete failure at 40 Hz was observed at and above 32°C in some preparations. Representative traces from a single trial showing the effects of temperature on muscle contraction are shown in Figure 6B. Calculating the Q10 within the negatively linear component of the graph (between 17°C and 27°C) generates a value of 0.24, accentuating this negative thermal dependence. Earlier experiments revealed that greater plasticity in muscle performance was observed when stimulating between 20 and 30 Hz. Consequently, we repeated this experiment using 25 Hz to determine if temperature might have a more profound effect, particularly at lower temperature. Significant differences were observed at 15, 16, 19, and 20°C, indicating more substantial effects of temperature at lower stimulation frequencies (Figure 6C). Rearing animals at different temperatures, particularly ectothermic animals, has been shown to adapt various aspect of an animal’s physiology, shifting their physiological efficiency closer to rearing temperatures (Bennett, 1985). Thus, we reared animals at 22, 25 and 29°C to determine if muscle performance could be shifted rightward towards better performance at higher temperatures. No significant difference was observed in animals reared at distinct temperatures (Figure 6D), suggesting the muscle excitation-contraction machinery does not display temperature adaptation.

**Figure 6.**
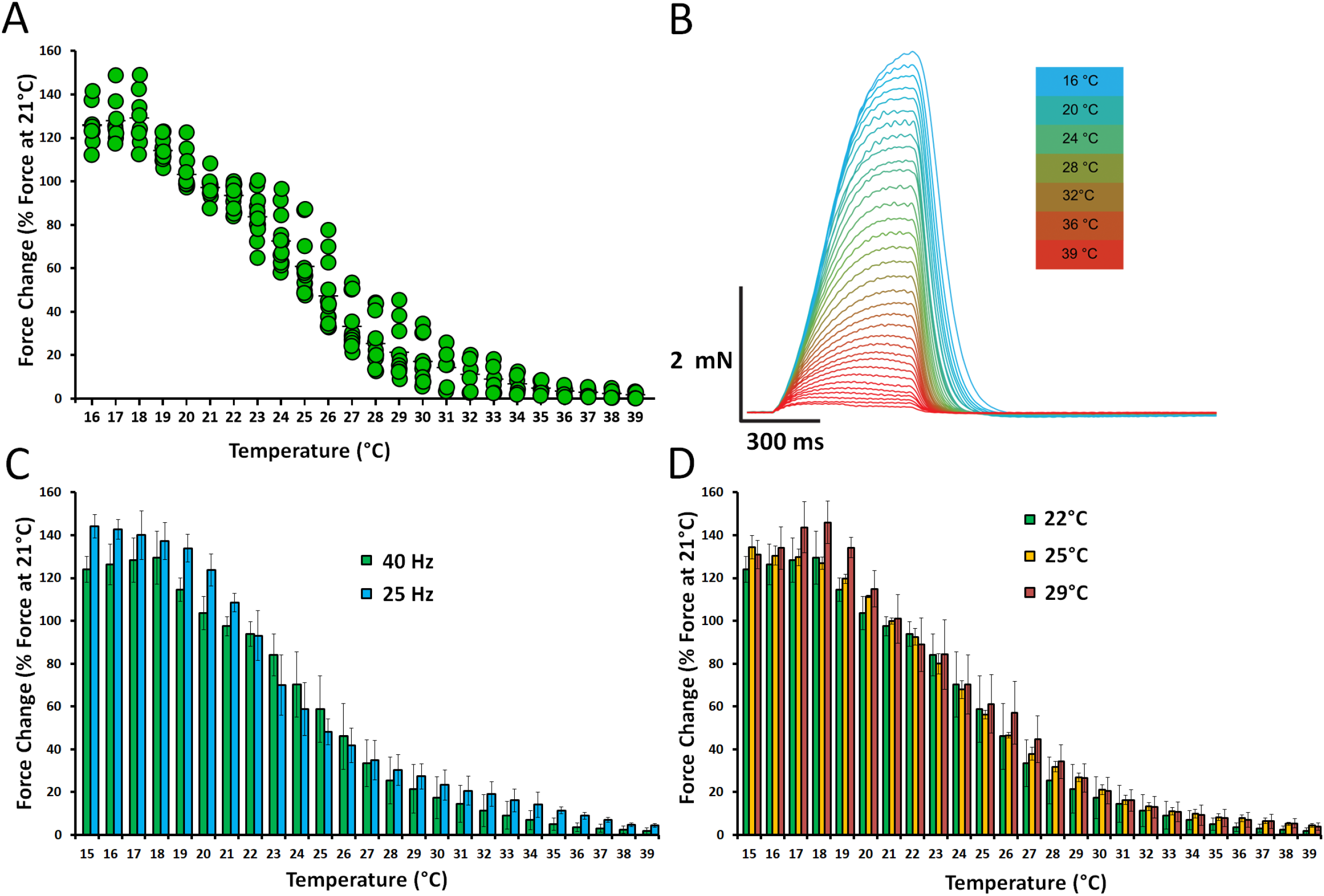
Effects of temperature on muscle contraction force. **A)** Force of muscle contraction as a function of temperature. Contractions were elicited using a 40 Hz 600 ms duration stimulus given continuously throughout the experiment as temperature was varied. **B)** Color-coded individual traces from a representative experiment depicting the effects of temperature on contraction amplitude. **C)** Histogram showing the effect of temperature on muscle contractions elicited using 25 Hz or 40 Hz stimulation for 600 ms duration. **D)** Effects of thermal acclimation on muscle contraction force assayed in larvae from stocks reared for 3 generations at 22, 25, or 29°C.

To examine how synaptic dysfunction might alter muscle contractile force, force-frequency plots were generated for several mutations that impair presynaptic neurotransmitter release or synapse number in Drosophila larvae. The magnitude of muscle force production elicited by 150 Hz stimulation for 600 ms (Figure 7A) or in response to a single stimulation (Figure 7B) was determined for several distinct synaptic mutants. Mutations in the SNARE-binding protein CPX significantly reduce evoked synchronous synaptic vesicle release and dramatically enhance spontaneous mini release (Cho et al., 2015; Huntwork and Littleton, 2007; Jorquera et al., 2012). Consistent with a key role for synchronous synaptic vesicle release for contraction force, *cpx* null mutant larvae show a significant reduction in both maximal muscle contraction force and the force elicited from a single stimulus (Figure 7A, B). Mutations in Synaptotagmin 1 (SYT1), a presynaptic Ca^2+^ sensor for synaptic vesicle fusion, eliminates synchronous evoked release and enhances asynchronous fusion (Guan et al., 2017; Lee et al., 2013; Yoshihara and Littleton, 2002). The maximal force produced in *syt1* null larvae was significantly reduced, even more substantially than in *cpx* mutants and consistent with the more severe defect in synchronous fusion in the absence of SYT1 (Figure 7A). Single stimuli were unable to generate contractions in *syt1* null larvae (Figure 7B), indicating muscle contraction in the absence of SYT1 likely requires the enhanced release from presynaptic facilitation that is intact in these mutants (Guan et al., 2020). SYT4 is a SYT isoform that localizes to the postsynaptic NMJ compartment and regulates retrograde signaling to enhance short-term plasticity following high frequency stimulation (Barber et al., 2009; Harris et al., 2016; Yoshihara et al., 2005). Loss of SYT4 caused a significant reduction in maximum force generation after a high frequency stimulation, but did not disrupt muscle contraction force elicited by single stimuli (Figure 7A, B). The *Drosophila* Gbb protein, a bone morphogenic protein (BMP) homolog, regulates synaptic growth, with null mutations reducing the number of synaptic boutons and release sites at the larval NMJ (McCabe et al, 2003). *Gbb* null mutants showed a significant reduction in both maximal force production and single stimulus contraction force. Together, these data indicate both synaptic release properties and synapse number contribute to excitation-contraction coupling in Drosophila larvae (Figure 7A, B).

**Figure 7.**
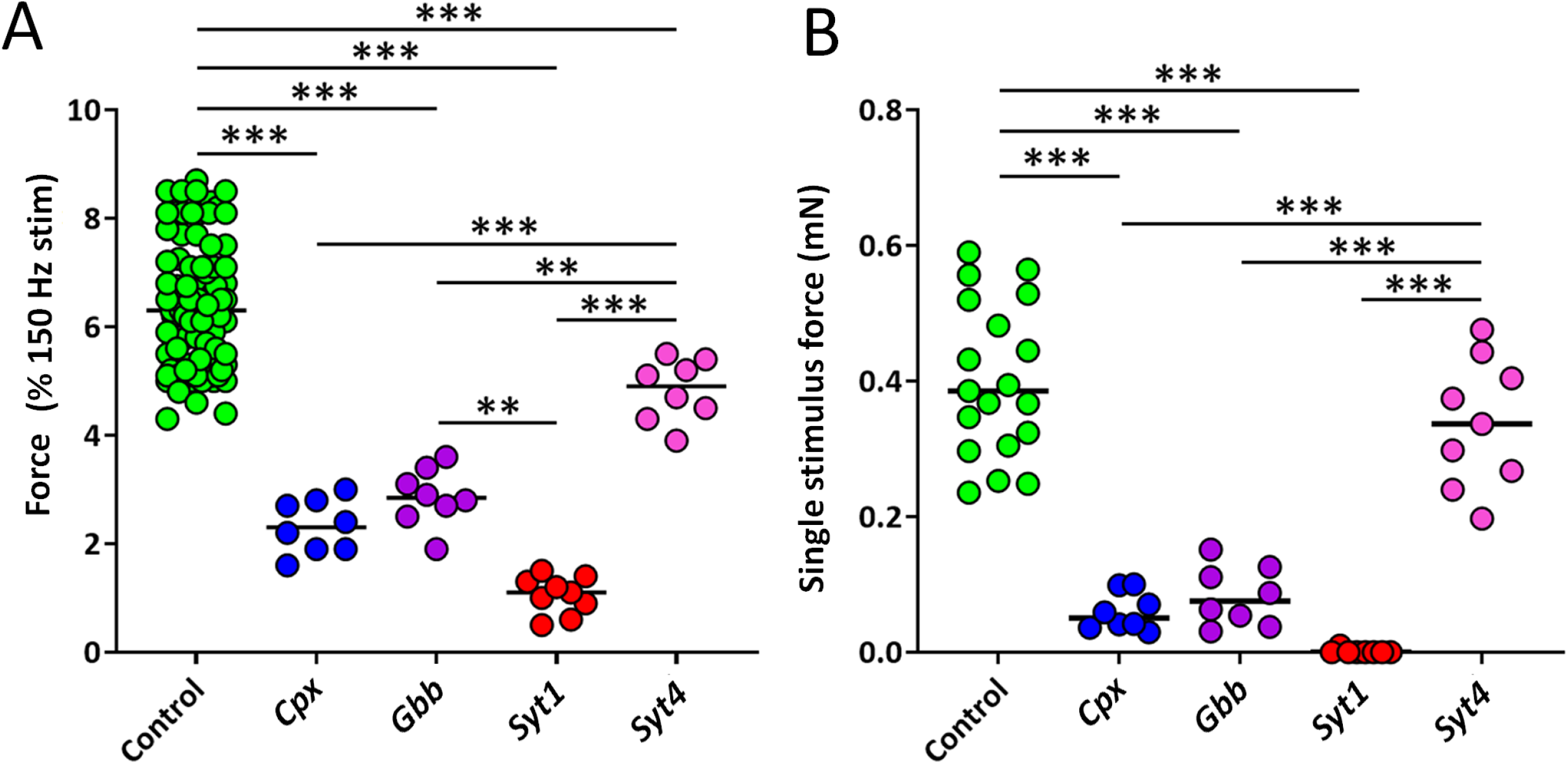
Effects of synaptic mutants on muscle contractile force. **A)** Maximal contraction force generated for control Canton S, *Cpx* null mutants (*cpx^SH1^*), *Gbb* null mutants (*Gbb^1^* /*Gbb^2^*), *Syt1* null mutants *(Syt1^ΛD4^/Syt1^N13^)* and *Syt4* null mutants (*Syt4BA^1^*). **B)** The magnitude of force generated from a single stimulus from each genotype is shown. One-way ANOVA, **p<0.01, ***p<0.001.

To explore pathways that regulate excitation-contraction coupling at the Drosophila NMJ, we assayed several potential modulators to examine their effects on muscle force production. Classical neuromodulators and neurotransmitters shown to affect muscle performance in other systems or in Drosophila were assayed. To initially screen these neuromodulators, a paradigm using steady-state 40 Hz stimulation for 600 ms with an interburst duration of 15 s was used as this stimulation generates robust muscle contractions but is below force saturation. Candidate neuromodulators were perfused onto larval preparations for 5 mins at the relatively high concentration of 10^-5^ M or 10^-6^ M to make sure any potential effects on contraction could be detected. Several neuromodulators failed to show any effect on steady-state muscle contraction force during stimulation, including 10^-5^ M leucokinin, 10^-5^ M gammaaminobutyric acid (GABA), 10^-5^ M acetylcholine, 10^-5^ M pituitary adenylate cyclase activating polypeptide (PACAP27), 10^-5^ M histamine, and 10^-5^ M dopamine (Figure 8A-F). In contrast, the TPAEDFMRFa peptide encoded by the *FMRFa* gene previously implicated in muscle performance (Hewes et al, 1998) robustly enhanced muscle contraction force. Exogenous application of 10^-6^ M TPAEDFMRFa steadily increased the amplitude of muscle contractions, reaching a maximal effect after 5 min of perfusion that resulted in an average force increase of 29 + 6.4% (Figure 8G). A dose-response curve for enhanced muscle contractile force was generated by examining the effects of TPAEDFMRFa at 10^-5^ M, 10^-6^ M, 10^-7^ M, 10^-8^ M, 10^-9^ M, and 10^-10^ M with 7 replicate animals for each concentration (Figure 8H). Under these conditions, the [EC_50_] of TPAEDFMRFa for enhancing muscle contraction was 5.38 x 10^-8^ M (Figure 8H).

**Figure 8.**
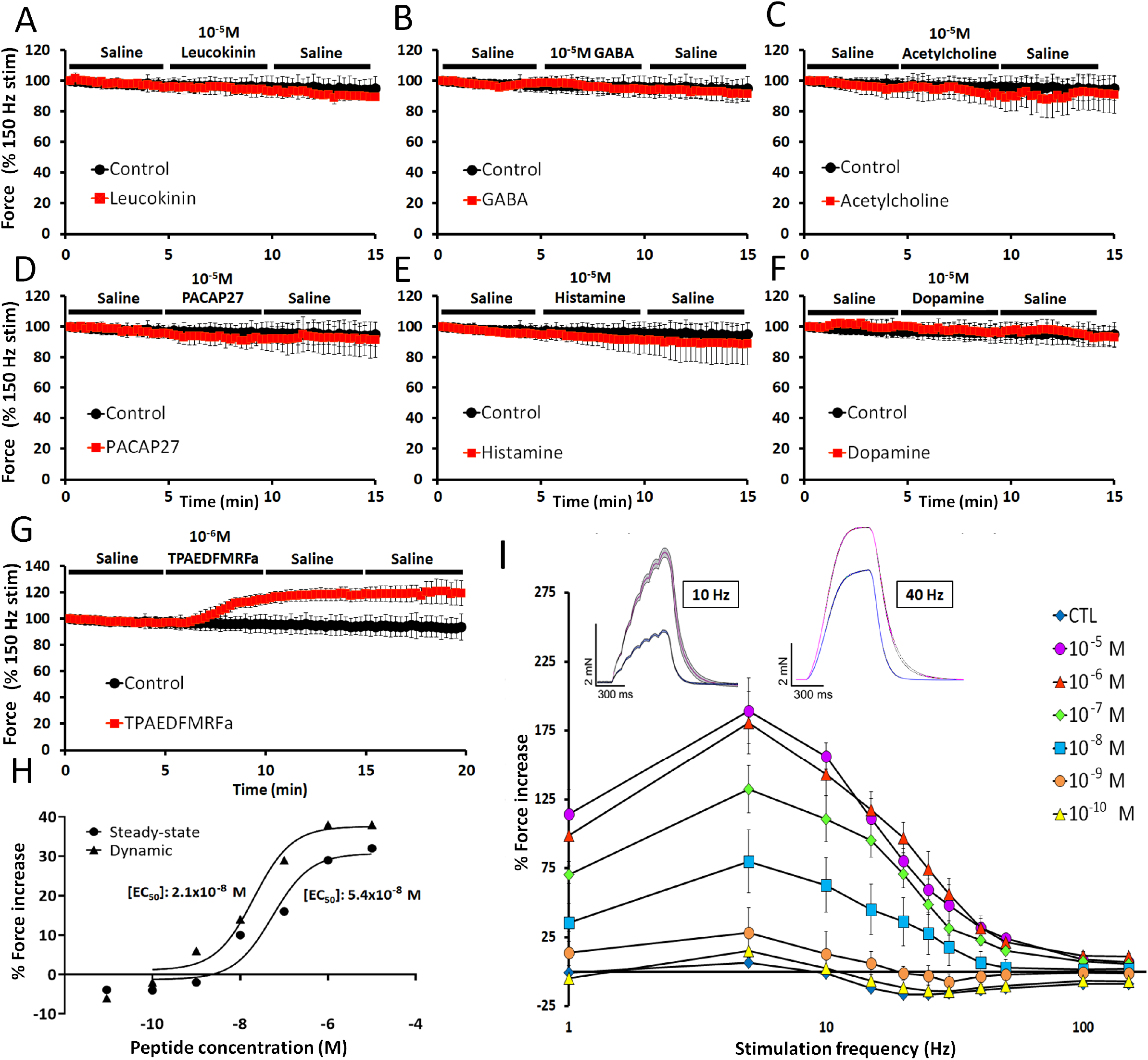
Amplitude of larval contractions plotted as a function of time using a steady-state motoneuron stimulation at 40 Hz for 600 ms duration every 15s. Experiments were 15-20 minutes in duration with a constant perfusion of saline for 5 minutes followed by addition of **A)** Leucokinin, **B)** GABA, **C)** acetylcholine, **D)** PACAP27, **E)** histamine, **F)** dopamine, or **G)** TPAEDFMRFa, followed by a 5-10 min saline washout. **H)** Dose-dependent effects of TPAEDFMRFa on muscle contraction using either the steady-state or dynamic motoneuron stimulation paradigm. **I)** Dose-dependent effects of TPAEDFMRFa using the dynamic motoneuron stimulation paradigm. Inset: control (blue) vs. TPAEDFMRFa (pink) at 10 Hz and 40 Hz stimulation.

To further characterize this neuromodulator, the same series of TPAEDFMRFa concentrations were applied using a dynamic force-frequency curve paradigm. Control recordings were conducted by running larvae through the stimulation protocol from 1-150 Hz, waiting a period of 5 min, then performing the same stimulation protocol a 2^nd^ time. For experimental trials, the neuropeptide was perfused in during the 5 min rest period between trials and throughout the entirety of the 2^nd^ stimulation period. The effects of TPAEDFMRFa on muscle force across each frequency and at each concentration are summarized in Figure 8I. At 40 Hz, the [EC_50_] was 2.1×10^-8^ M using the dynamic protocol (Figure 8H). There were no statistical differences between TPAEDFMRFa concentrations at 100 and 150 Hz stimulation, although significant increases were observed at 50 Hz stimulation from 10^-7^ to 10^-5^ M. Surprisingly, stimulation frequency decreases contraction efficiency as TPAEDFMRFa concentration increases, peaking at 5 Hz where a nearly 200% increase in contraction force is observed at 10^-5^ and 10^-6^ M. Based on control conditions (Figure 2), we predicted that the peptide would exert the greatest effect between stimulation frequencies of 20 to 35 Hz where the contractile machinery is normally most sensitive to nerve output. In contrast, the effectiveness of the peptide increased as stimulation frequency decreased. EC_50_ values generated for each stimulation frequency revealed similar values in the 2.1 to 3.9 x 10^-8^ M range (supplemental table 1).

To begin elucidating the molecular pathway by which TPAEDFMRFa exerts its effects on contraction force, the UAS-GAL4 system was employed to knockdown the FMRFa receptor using UAS-RNAi against *FMRFa* mRNA. The pan-neuronal driver *Elav*-GAL4 was used to express *FMRFa* RNAi presynaptically in motoneurons and the muscle specific *Mef2*-GAL4 was used to drive the RNAi postsynaptically in muscles. Knocking down the FMRFa receptor either pre- or post-synaptically significantly reduced the ability of TPAEDFMRFa to enhance muscle contraction force (Figure 9A). Using both GAl4 drivers to simultaneously knockdown the FMRFa receptor pre- and post-synaptically resulted in an additive decrease in the ability of the peptide to enhance muscle contraction. However, TPAEDFMRFa was still able to partially increase contraction even when the FMRFa receptor was knocked down simultaneously with both drivers. In addition to the FMRFa receptor, myosuppressin receptors have also been shown to mediate some behavioral effects for a different FMRFa peptide, DPKQDFMRFa (Klose et al. 2010). To determine if TPAEDFMRFa might also activate this pathway, the two myosuppressin receptors encoded in the genome were targeted with independent RNAi lines to assay their involvement in TPAEDFMRF’s ability to enhance muscle contraction. Pre- and post-synaptic expression of RNAi against the Dromyosuppressin receptor 1 did not significantly reduce the ability of the peptide to potentiate muscle contraction (Figure 9B). However, knockdown of Dromyosuppressin receptor 2 with either *Mef2-GAL4* or with both *Mef2-GAL4* and *Elav-GAL4* simultaneously significantly reduced the ability of TPAEDFMRFa to potentiate muscle contraction force. Taken together, these results indicate the TPAEDFMRFa neuropeptide enhances muscle contractions through activity in both pre- and post-synaptic compartments via the FMRFa receptor and Dromyosuppressin receptor 2.

**Figure 9.**
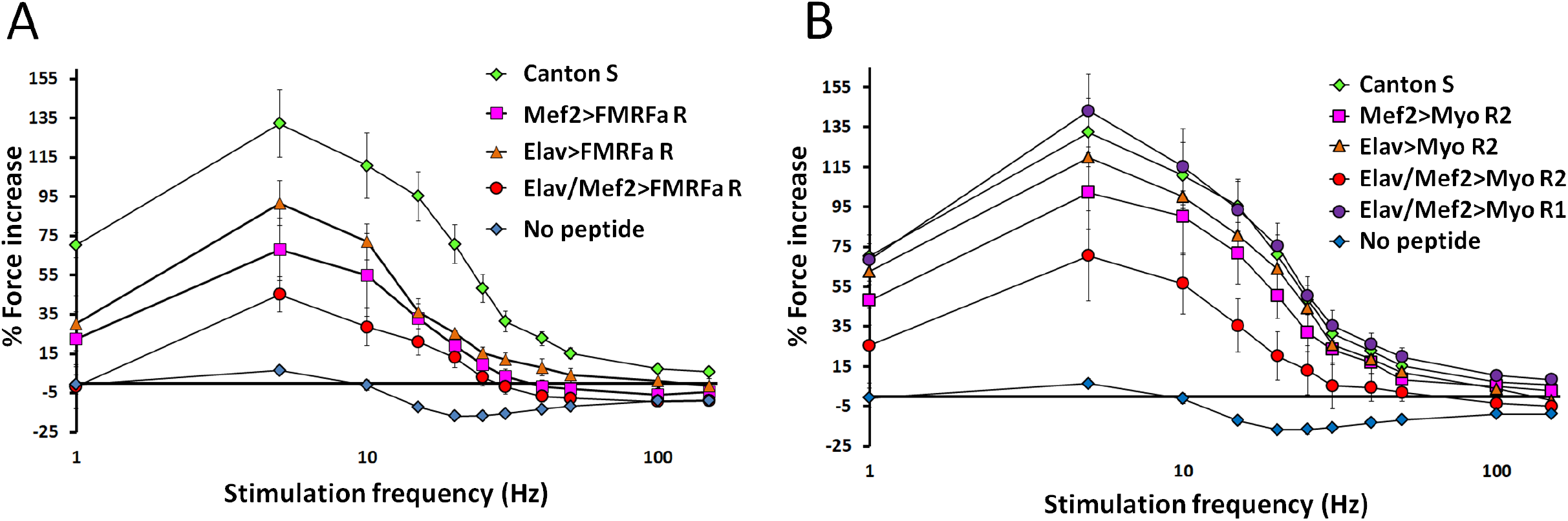
Analysis of the TPAEDFMRFa pathway for enhancing contractile force at the NMJ. A) Force-frequency plots for the effectiveness of TPAEDFMRFa on muscle contraction force in Canton S versus larvae expressing FMRFa receptor UAS-RNAi presynaptically (Elav-GAL4), postsynaptically (Mef2-GAL4) or in both compartments. B) Force-frequency plots for the effectiveness of TPAEDFMRFa on muscle contraction force in Canton S versus larvae expressing Myosuppressin receptor 1 or 2 UAS-RNAi presynaptically (Elav-GAL4), postsynaptically (Mef2-GAL4) or in both compartments.

## Discussion

These data demonstrate that the excitation-contraction coupling machinery in *Drosophila* 3^rd^ instar larvae provides a valuable model for exploring and dissecting the component parts that contribute to neuromotor circuitry. Using a force transducer with 10 μN resolution, we assayed the contribution of key biophysical, physiological, genetic and molecular parameters to muscle contractility. The force-frequency relationship in bodywall muscles was determined using a dynamic motoneuron stimulation paradigm to quantify the full range of muscle contractility, from threshold to saturation. Larval size and orientation did not influence muscle performance, although physiological saline composition, predominately magnesium and calcium concentrations, had a profound effect on muscle contractility. Several mutants that disrupt synchronous synaptic vesicle fusion, including *Syt1* and *Cpx*, dramatically reduced the magnitude of muscle contractions over all stimulation frequencies. In addition, mutations in *Gbb* that reduce synapse number also decreased the overall force of contraction, suggesting both the strength of synaptic transmission and the number of release sites are important for generating muscle contractile force. Changes in temperature also profoundly impacted muscle performance, with subtle decreases in temperature leading to a 30% increase in muscle force. Increasing the temperature beyond 30°C resulted in a near complete loss of muscle force production. Many different conventional invertebrate neurotransmitter and neuromodulators were shown to have no impact on muscle performance. However, exogenous application of the neuropeptide TPAEDFMRFa led to a dose-dependent increase in muscle force by as much as 300%. Genetically reducing the expression of the FMRFa and Myosuppressin 2 receptors significantly attenuated the potency of the peptide. Both pre- and post-synaptic mechanisms contributed to TPAEDFMRFa’s capacity to enhance contraction force, suggesting it acts through multiple receptors and in both synaptic compartments to effect muscle output.

Our data indicate neither the size of 3^rd^ instar larvae nor the muscle subgroup examined had any significant effect on the maximal force production or biophysical properties of excitation-contraction coupling across the whole animal (Figures 3, 4). Although muscle architecture can impact force production (Burkholder et al., 1994; David et al., 2016), studies have shown that muscle length does not result in greater force production, but rather impacts contraction velocity (Forman et al., 1972). For some fibers, e.g. muscle fiber 6, a clear increase in the resting muscle width is observed, compared to its direct neighbor, muscle fiber 7. It is interesting that a greater number of longitudinal muscles are likely present along the ventral axis of the larvae (Fig. 4, e.g. 6, 7, 12, 13) compared to the dorsal axis (1, 2, 9, 10). Additionally, considerably more muscles lie along the ventral axis that are not directly longitudinal, but have insertion angles that would generate considerable forces along the longitudinal axis (e.g. 14, 15, 16, 17, 26, 27, 28, 30) compared to the dorsal axis (3, 11, 19, 20). Thus, given substantially more muscle fibers contribute to longitudinal force production, it is possible that dorsally located muscles produce greater force.

Changes in external Ca^2+^ have a profound effect on neurotransmitter release, as release is proportional to the third to fourth power of Ca^2+^ entry (Augustine et al., 1987; Hille, 1985; Katz and Miledi, 1970). Increases in the concentration of external Ca^2+^ at the Drosophila NMJ have a sigmoidal relationship with excitatory junctional currents, saturating at ~1.0 mM (Roche et al., 2002; Rohrbough et al., 1999). Figure 5E demonstrates the dose-dependent effects of external Ca^2+^ on EJP amplitude, which saturate at ~1.0 mM. Increasing external Ca^2+^ has nearly identical effects on muscle force production as on synaptic properties observed from electrophysiological recordings, showing a sigmoidal curve which saturates with 1.0 mM external Ca^2+^ in Drosophila HL3.1 saline. This effect is also conserved when examining the force of contractions elicited from a single action potential in HL3.1 saline. Thus, the effects of external Ca^2+^ on synaptic physiology appear to couple in a 1:1 fashion with the contraction machinery.

The divalent cation magnesium (Mg^2+^) readily moves throughs Ca^2+-^channels and therefore competitively interacts with Ca^2+^ entry. Mg^2+^ also has greater electronegatively than Ca^2+^ and can directly block Ca^2+^ channels (Kuno and Takahashi, 1986). In *Drosophila*, Jan and Jan (1976) reported a significant change in quantal content following a 2 mM shift in Mg^2+^ concentration in standard saline (from 2 to 4 mM), and similar effects were observed in mouse vas deferens (Bennett and Florin, 1975). We observed significant effects when increasing the external Mg^2+^ concentration from 4 mM to 20 mM. At low external Ca^2+^ concentrations (0.1 mM), contractions were not observed at any stimulation frequency or duration with 20 mM Mg^2+^, compared to robust contractions observed in 4 mM Mg^2+^ (Fig 5A). A significant decrease in muscle force production was apparent in 0.25 and 0.5 mM [Ca^2+^]_o_ in 4 mM vs. 20 mM [Mg^2+^]_o_. Indeed, a significant difference was observed in the amplitude of contractions elicited from single action potentials at all Ca^2+^ concentrations investigated between the two Drosophila salines (Fig 5C-D). While increasing external Mg^2+^ concentration has been shown to decrease synaptic strength, Mg^2+^ is also known to have a strong effect on muscle cells. Current-clamp recordings from L-type Ca^2+^ channels, similar to those found in *Drosophila* muscle cells, reveal a significant reduction in Ca^2+^ current following modest increases in external Mg^2+^ concentrations (Wang et al., 2004). Additionally, Mg^2+^ has been shown to impact the SR Ca^2+^-content (Launikonis and Stephenson, 2000), efficiency of the Na^+^/ Ca^2+^ exchanger (Levitsky and Takahashi, 2013), as well as the uptake efficiency of the Ca^2+^ ATPase pump located on the SR (Michailova et al., 2004). Thus, the observed effects of Mg^2+^ may be a consequence of presynaptic effects on Ca^2+^ channels, or numerous effects directly on muscle fiber machinery.

Feng et al. (2004) noted a gradual but significant reduction in the EJP amplitude with increases in temperature, ultimately resulting in the inability to elicit an EJP at 40-42°C. Our results demonstrate that increasing temperature also gradually and significantly decreases the force of muscle contraction, ultimately resulting in the inability to elicit contraction between 34 and 39°C (Fig. 7). Action potential propagation in axons is known to be blocked at higher temperatures due to rapid gating dynamics of Na^+^ channels, reducing positive charge influx during membrane depolarization (Westerfield et al, 1978). Changes in temperature also have a substantial effect on the amplitude and waveform of action potentials, with decreases in temperature increasing the amplitude of action potentials, and drastically slowing the rise time (τ-rise) and decay time (τ-decay) (Miller and Rinzel, 1981). Impulse propagation failure is thought to occur at regions of nonuniform morphology causing increases in capacitance, typical at axonal branch points and transitions zones from myelination to demyelination (Hille, 1985). Temperature and conduction velocity of APs in axons and muscles however are positively correlated (Miller and Rinzel, 1981). The temperature-dependent effects on EJP amplitude observed by Feng et al., (2004) appear to strongly correlate with the effects of temperature on muscle contraction dynamics observed here (Figure 7). Our results also found that the strength of muscle contractions increases with decreases in temperature, saturating at 17-18°C. In addition, to the effects on axonal conduction properties, temperature may also affect multiple aspects of muscle contraction from membrane excitation (Vornanen, 2016), Ca^2+-^dynamics from the SR (Abu-Amra et al., 2015; Shiels et al., 2002), Ca^2+^-dependent contraction activation (Ranatunga, 2018), and mechanisms of cross-bridge cycling (Wang and Kawai, 2001). While effects of temperature on synaptic efficacy appear to directly translate to the excitation-contraction coupling machinery, it is possible that temperature is also affecting multiple aspects of muscle biology.

To examine how defects in synaptic transmission and synapse number manifest onto excitation-contraction coupling, we quantified the force-frequency relationship for mutations in *Syt1, Cpx, Gbb* and *Syt4*. Loss of the Ca^2+^ sensor SYT1 caused a 100% and 84% reduction in the amplitude of single and maximal contractions, respectively, compared to a 90% reduction in EJC amplitude (Guan et al, 2017). Loss of CPX resulted in a 85% and 63% reduction in the amplitude of single and maximal contractions compared to a 75% reduction observed in EJP amplitude (Cho et al, 2015). These results indicate the effects of these mutations on evoked synaptic transmission relate directly to effects on the contraction machinery. Mutations in Syt4, which regulates retrograde signaling and short-term synaptic plasticity, caused a significant 24% decrease in the force of maximal contractions, but no effect on the amplitude of contractions elicited from single action potentials. A similar 20% reduction in EJC amplitude was also found in *Syt4* mutants (Barber et al, 2009). *Gbb* mutants displayed a 80% and 55% decrease in contraction force for single stimuli and maximal contraction. Disruption of GBB signaling leads to a severe decrease in synapse number and a corresponding 70% decrease in EJP amplitude (McCabe et al, 2003). Overall, these data indicate whole larval bodywall force contraction assays provide a robust readout for synaptic dysfunction, with single contraction data closely matching previously reported defects in evoked release.

The NMJ of arthropods have long been explored as critical areas of influence by neuromodulatory substances (Robbins, 1959; Van Harreveld and Mendelson, 1959). Among the small classical neuromodulators, we assayed muscle contraction force changes following bath application of dopamine, histamine, acetylcholine and GABA. The biogenic amine dopamine has been shown to enhance synaptic efficacy and muscle contraction in numerous model systems, including the lobster (Lingle, 1981), Aplysia (Swann et al., 1982) and shrimp (Meyrand and Moulins, 1986). Additionally, dopamine has previously been shown to decrease EJP amplitude at Drosophila larval muscle 6 (Cooper and Neckameyer, 1999). However, we found no observable effect of dopamine on muscle contractions. Perhaps only subsets of muscle fibers are affected by dopamine, and the altered EJPs in these fibers is insufficient to generate a noticeable enhancement in muscle force production over the entire larval bodywall. Although histamine has been shown in modulate the force of contraction of smooth muscle and cardiac tissue (Chan, 1977; Reite, 1972), we found no effect on the Drosophila NMJ. Acetylcholine functions as the neurotransmitter at vertebrate NMJs, but can also serve as a neuromodulator (Picciotto et al., 2012). Exogenous application of acetylcholine had no effect on contraction force at the Drosophila glutamatergic NMJ. GABA is classically defined as the primary inhibitory neurotransmitter in the brain, but no evidence exists to suggest GABA receptors are present at the NMJ (Kravitz et al., 1963). Indeed, GABA had no effect on modulating force at the Drosophila NMJ. These observations suggest glutamate is likely to be the major small molecular effector at the fly NMJ.

Multiple neuropeptides have also been suggested to modulate the neuromuscular system. Landgraf et al, (2003) described leucokinin-immunoreactivity in type III terminals at the Drosophila NMJ, but we found no effect on contraction when leucokinin was exogenously applied to the dissected larval preparation. Pituitary adenylate cyclase activating polypeptide (PACAP) is encoded by the amnesiac gene in Drosophila. PACAP-38 has previously been shown to modulate Ca^2+^ channel dynamics and contraction in Drosophila 3^rd^ instar bodywall muscles (Bhattacharya et al., 2004; Zhong and Peña, 1995). We found that the shorter isoform, PACAP-27, had no effect on muscle contraction. The *Drosophila* genome encodes a single gene for the FMRF-like peptides, the *dFMRFamide* gene which encodes 8 peptides. DPKQDFMRFa is the most abundantly encoded, with 5 replicate copies, and has been previously characterized at the Drosophila NMJ. DPKQDFMRFa is known to enhance synaptic transmission via the CaMKII pathway through the FMRFa receptor (Dunn and Mercier, 2005; Klose et al, 2010). DPKQDFMRFa increases nerve-evoked contractions and can also elicit contractions in the absence of the CNS (Hewes et al, 1998; Clark et al, 2008; Ormerod et al, 2015).

The second most abundant FMRFa-like peptide is TPAEDFMRFa with two replicate copies (Nambu et al, 1988; Schneider and Taghert, 1988). Previous work has shown that TPAEDFMRFa enhances twitch tension in a reduced preparation from Drosophila 3^rd^ instar larvae in a similar magnitude to DPKQDFMRFa (Hewes et al, 1998). To further examine this pathway for modulating neuromuscular contraction, we examined the effects of exogenous application of TPAEDFMRFa by eliciting contractions using 40 Hz stimulation. This paradigm revealed a robust increase in the amplitude of nerve-evoked contractions that did not wash out with 10 min of saline perfusion (Figure 8G). TPAEDFMRFa displayed an EC_50_ of 5.4 x 10^-8^M, generating a 20% increase in the amplitude of contractions to a maximal increase of 38% at 10^-6^M. By examining the effects of the peptide across a wide-range of stimulation frequencies from 1-150 Hz, TPAEDFMRF was much more effective at increasing contraction force at lower stimulation frequencies, with maximal effects observed at 5 and 10 Hz (189 ± 28% and 156 ± 10% respectively). Given each successive decrease in stimulation frequency elicited a greater amount of potentiation by the peptide, it is likely that the electrochemical driving force and biophysical properties inherent to the contractile machinery have the greatest potential for modulation at lower stimulation frequencies given they are the farthest from saturated. The lack of modulation at higher stimulation frequencies is likely a result of muscle contraction being at or near saturation. Thus, release of neuromodulatory substances like FMRFa peptides would maximize the amplitude of contractions, enabling larvae to generate a stronger contraction with less synaptic input. Release of the peptide could aid in the fight-or-flight response to enable larvae to reach maximal locomotor velocity or play a role in preventing synaptic fatigue under conditions of chronic neuromotor activation.

Using the UAS/GAL4 system to knockdown putative receptors mediating the effects of TPAEDFMRFa, we found a role for the FMRFa receptor in both pre-and postsynaptic compartments contributes to the peptide’s ability to potentiate nerve-evoked contractions. However, using nerve and muscle-specific drivers simultaneously was insufficient to completely abolish the effect on contraction force. It is therefore likely that other receptors also contribute to the enhancement of nerve-evoked contractions by the peptide, or RNAi was insufficient to fully remove the FMRFa receptor. We tested knockdown of two myosuppressin receptors previously implicated in mediating the effects of DPKQDFMRFa (Klose et al, 2010). Myosuppressin receptor 1 appears to play no role in the enhancement of nerve-evoked contractions by TPAEDFMRFa. However, knocking down myosuppressin receptor 2 either postsynaptically, or pre and post-synaptically, significantly reduced the ability of the peptide to enhance contraction.

Taken together, our results demonstrate that Drosophila serves as a robust system to explore excitation-contraction coupling and dissect the component parts that contribute to neuromotor circuitry. Our characterization of the force-frequency components of contraction indicate Drosophila muscle contractile force is similar to force-frequency data obtained in other vertebrate models (Eshima et al., 2017; Terry et al., 2014). Indeed, critical muscle genes underlying the development and function of the musculature are highly conserved between Drosophila and more commonly studied vertebrate species (Taylor, 2013). As such, Drosophila larval bodywall muscles provide an excellent model for investigations of excitation-contraction coupling and linking known synaptic defects in neuronal mutants to their final effect on muscle force production.

## Materials and methods

### Drosophila stocks

*Drosophila melanogaster* were cultured on standard medium at 21°C at constant humidity in a 12:12 light:dark cycle. Wandering 3^rd^ instar larvae of both sexes were used for experimentation. Canton S (CS) flies obtained from the Bloomington Drosophila stock center (BDSC) were used as controls except when noted. *Gbb* mutants (*Gbb^1^*/*Gbb^2^*) was obtained from BDSC (McCabe et al., 2003). The remaining mutants examined included *Cpx* nulls (Huntwork and Littleton, 2007), *Syt1* nulls (*Syt1^N13^/Syt1AD^4^*) (Littleton et al., 1994) and *Syt4* nulls (*Syt4^BA1^*) (Yoshihara et al., 2005). Other genotypes used in the study include: *elav^C155^*-GAL4 (BDSC #8765), *Mhc*-GAL4 (BDSC#55132), FMRFa receptor RNAi (VDRC v9594), Dromyosuppressin receptor 1 RNA (VDRC v9369), Dromyosuppressin receptor 2 (VDRC v49952).

### Dissection

Wandering 3^rd^ instar larvae were isolated from the sides of culture vials and dissected in modified hemolymph-like (HL3) saline. Two *Drosophila* salines were examined (Feng et al., 2004). The first Hemolymph-Like saline HL3 has the following composition (in mM): NaCl: 70; KCl: 5; CaCl_2_:1.5; MgCl_2_: 20; NaHCO_3_: 10; Trehalose: 5; Sucrose: 115; HEPES: 5, (pH = 7.18). The second saline, HL3.1, has the same composition but [MgCl_2_] was reduced to 4 mM. Larvae were pinned dorsal side up at the anterior and posterior ends, a small incision was made along the entire dorsal midline and the visceral organs were removed. All nerves emerging from the central nervous system (CNS) were severed at the ventral nerve cord, and the CNS and ventral nerve cord were removed. In some experiments, the orientation of the larvae prior to initial dissection was altered depending on which muscles were examined. To examine the dorsal muscles, the animal was pinned ventral side up and dissected along the ventral midline. To examine the lateral muscles, the animal was pinned lateral side up (muscle fiber 4 at the apex) and dissected along the lateral most point.

### Electrophysiology

Excitatory junctional potentials (EJPs) were elicited by stimulating severed abdominal nerves. A Master 8 A.M.P.I. (Jerusalem, Israel) stimulator was used to elicit stimulation via a suction electrode (A-M systems, Sequim, WA). EJPs were recorded using sharp glass microelectrodes containing a 2:1 mixture of 3M potassium chloride:3 M potassium acetate, with an electrode resistance of 40-80 MΩ. An Axoclamp 2B amplifier (Molecular Devices, San Jose, CA) was used for signal detection and digitized via Axon Instruments digidata 1550 (Molecular Devices). Signals were acquired at 10 kHz using Clampex and processed using Clampfit and MiniAnalysis.

### Nerve-evoked Contraction Force Recordings

All force recordings were obtained using the Aurora Scientific 403A force transducer system (Aurora Scientific, Aurora, Canada), including the force transducer headstage, an amplifier, and digitizer. Nerve-evoked contractions were generated using bursts of electrical stimuli from a Master 8 (A.M.P.I.) stimulator. The duration of single impulses was 5×10^-4^ s and the interburst duration was kept constant at 15 s. Burst duration and frequency were altered for each individual experiment. Larvae were dissected as outlined above. To attach larvae to the force transducer, a hook was made from a fine minuten pin and placed onto the posterior end of the larvae. Digitized data was acquired using Aurora Scientific software, Dynamic Muscle Acquisition Software (DMCv5.5). The digitized data was imported and processed in Matlab using custom code written by A. Scibelli (available on request). Temperature was controlled using a CL-200A heater-cooler controller (Warner Instruments, Hamden CT) with a heat sink (Koolance) that regulates temperature via feedback from a thermocouple placed directly next to the animal to enable control within 0.1 °C. Temperature was rapidly modified by constant perfusion of physiological saline though a Harvard apparatus in-line saline temperature controller (Holliston, MA) and a Warner Instruments dish temperature regulator (Model TB3 CCD).

### Pharmacological agents

Histamine hydrochloride (H7250, Sigma, St. Louis, MO), dopamine hydrochloride (H8502, Sigma), PACAP27 amide, Ovine (2151, Sigma), GABA (A2129, Sigma), Acetylcholine chloride (A6626, Sigma). Leucokinin (NSVVLGKKQRFHSWG) and TPAEDFMRFa were custom synthesized from Genscript (Piscataway, NJ) with a >98% purity.

## Acknowledgements

This work was supported by NIH grant NS40296 to J.T.L and a postdoctoral research fellowship from Natural Science and Engineering Research Council of Canada to KGO. We thank the Bloomington Drosophila Stock Center (NIH P40OD018537) for Drosophila stocks and members of the Littleton lab for helpful discussions and comments on the manuscript.

## Supplemental Table

**Supplemental Table 1:** EC_50_ calculations for TPAEDFMRFa’s effectiveness for increasing the force of contractions. For each frequency the maximum force increase (typically from 10^-5^ M) observed following TPAEDFMRFa was also calculated.

| [EC50] for TPAEDFMRFa |  | Maximum force increase (%) |
| --- | --- | --- |
| 1 | 3.90 x 10<sup>-8</sup> M | 115 |
| 5 | 2.60 x 10<sup>-8</sup> M | 183 |
| 10 | 1.85 x 10<sup>-8</sup> M | 156 |
| 15 | 1.41 x 10<sup>-8</sup> M | 130 |
| 20 | 1.13 x 10<sup>-8</sup> M | 114 |
| 25 | 1.23 x 10<sup>-8</sup> M | 91 |
| 30 | 1.27 x 10<sup>-8</sup> M | 72 |
| 40 | 2.10 x 10<sup>-8</sup> M | 39 |
| 50 | 2.43 x 10<sup>-8</sup> M | 34 |
| 100 | 1.30 x 10<sup>-8</sup> M | 21 |
| 150 | 6.1 x 10<sup>-9</sup> M | 20 |

## Supplemental figure legends

**Supplemental Figure 1 (S1):**
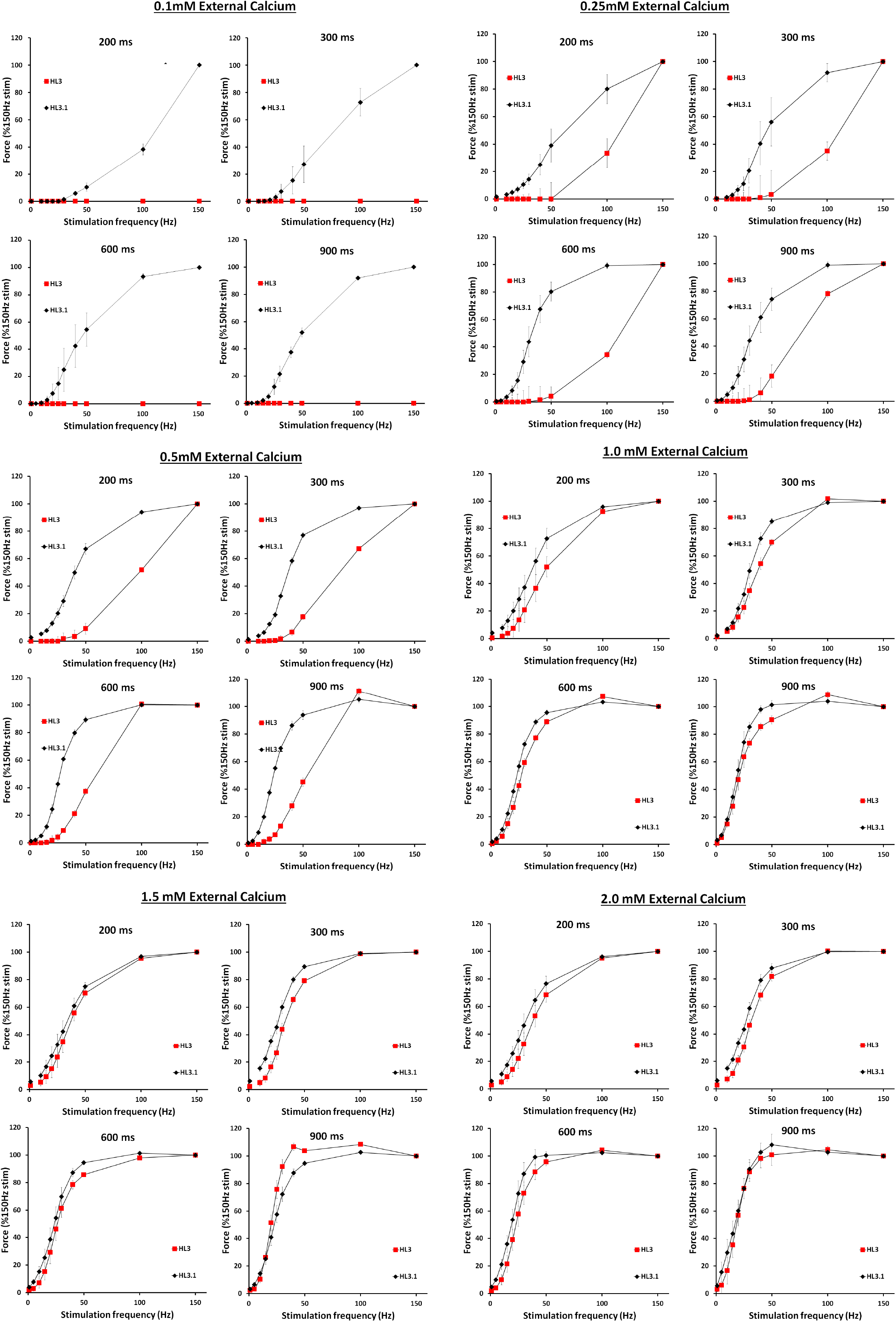
Effects of external Ca and Mg on muscle contraction force. Force-frequency curves generated using 200, 300, 600, and 900 ms duration stimuli for two physiological salines, HL3 (20 mM Mg^2+^) and HL3.1 (4 mM Mg^2+^) in 0.1 mM [Ca^2+^]_o_, 0.25 mM [Ca^2+^]_o_, 0.5 mM [Ca^2+^]_o_, 1.0 mM [Ca^2+^]_o_ and 2.0 mM [Ca^2+^]_o_.

